# Defusing the consumption bomb

**DOI:** 10.64898/2026.05.06.723357

**Authors:** Vanessa P. Weinberger, Ethan Duncil, Kimberly J. Cook, Miikka Tallavaara, Mikael A. Manninen, Jordan G. Okie, Trevor S. Fristoe, Joseph R. Burger

## Abstract

For decades, the “population bomb” has dominated environmental discourse, arguing that high fertility rates –especially in the low income countries– drive global environmental problems. However, current trajectories show global declines in population growth rates, especially in higher development index (HDI) nations, which have the highest consumption. Here we showcase evidence for a paradigm shift from the “population bomb” to a “consumption bomb” narrative of the Anthropocene emphasizing the central role of increases in per capita energy use and CO_2_ production, modulated by current standard metrics for development and affluence, in transforming the Earth system. Defusing and manoaging the consumption bomb requires rethinking economic growth and wellbeing metrics, reallocating resources toward global change retribution and mitigation, especially in low HDI countries, and transitioning from continually-increasing energy expenditures, especially from fossil fuels, toward more equitable and ecologically resilient ways of living. A new sustainability science must move beyond population counts to confront the biophysical and energetic consequences of the changing cultural, economic, and technological systems that sustain ever-growing demands on Earth’s life-support systems.

## Introduction

The 1968 publication of *The Population Bomb* by the late Paul and Anne Ehrlich sounded a global alarm: unchecked population growth would soon outpace food production and overwhelm ecological capacity, triggering famine, disease and conflict (Ehrlich 1968)^1^. This book echoed the Reverend Thomas Malthus almost 200 years earlier (Malthus 1798, Burger et al. 2019). Ehrlich’s book catalyzed decades of political and even “scientific” efforts to curb fertility rates, especially in low income countries, and cemented the “population bomb” as a cornerstone of environmental discourse for more than half a century. Although the world’s population has more than doubled since the publication of Ehrlich’s book, most countries are now approaching or are below replacement. Current projections suggest that we could approach Zero Population Growth (ZPG) in the coming decades. What continues to increase near-exponentially, however, is the human demand for Earth’s resources.

Following the population bomb, Ehrlich and Holdren (1971) implied the interacting roles of consumption and population size in driving environmental degradation in the IPAT equation (Impact = Population × Affluence × Technology). Yet popular discourse, academic discussion and even policy seemed to fixate on population size (“P”), overlooking the outsized influence of affluence (“A”) and technology (“T”), that have been recognized in recent decades. For example, Bradshaw et al. (2021) and Cafaro et al. (2022) argue that overpopulation is a primary driver of biodiversity loss; the *Overpopulation Project* (n.d.) calls for deliberate population reduction as its motivation; and many researchers have linked immigration restriction to environmental protection (e.g., Hurlbert 2011, Axelrod 2018). In contrast, Hughes et al. (2023) argue that “smaller human populations are neither a necessary nor sufficient condition for biodiversity conservation”, while Bluwstein et al. (2021) pose that “rather than distracting attention through neo-Malthusian tropes, scientists should help expose the structural causes and drivers of inequality, overproduction and overconsumption”, emphasizing capitalism and governance failures as more immediate drivers of biodiversity decline. Without addressing affluence and consumption, reaching Zero Population Growth (ZPG) alone cannot achieve sustainability.

While population growth has slowed since the 1960s, annihilation of the Earth’s resources continues to intensify. In this time, global energy use has more than quadrupled, reflecting transformations in economies, technologies, and cultures (Moses 2009, Delong et al. 2010, Hamilton 2012, Burger et al. 2019). Industrialization, urbanization, technological complexification and the spread of wealthy and ultra wealthy lifestyles have driven surging demand for mobility, construction, meat, and electronics. Increasing consumption levels have pushed the Earth to approach and even surpass planetary limits (Burger et al. 2012, Steffen et al. 2018, Richardson et al. 2023, Tian et al. 2025). As long as current consumption trends proceed (Box 1), increasing per capita energy and material use will continue fueling steep increases in environmental impact, even where population growth has stabilized.

Understanding the current socio-environmental predicament requires an appreciation for the (cultural) evolutionary history and trajectory of our species – from small networks of hunter-gatherer societies to industrial-agricultural-technological societies. The “population bomb” narrative fails to capture these realities. Trends of energy and environmental demand in modern human societies are better characterized as a “consumption bomb”—the expanding appetites for lifestyles that require increasing demands on the Earth system and is pushing the planet toward dangerous ecological and societal tipping points (Figure 1, Barnosky et al. 2012, Steffen et al. 2018, Søgaard-Jørgensen et al. 2024). Technological innovations can push against biophysical constraints and increase the (short-term) capacity of the Earth system to support the human system. Throughout much of their history, humans have responded with near continuous exponential population growth. Following the industrial revolution and especially since the 1950’s (the Great Acceleration), pressure on Earth’s support systems has been driven increasingly by rising per capita consumption. While increasing per capita consumption and associated metrics of development contribute to the demographic transition and decelerating population growth, the trajectory of total consumption continues to climb −, sustaining such dynamics ultimately requires further technological evolution and expansion to avoid population or economic collapse (Cohen 1995, Weinberger et al. 2017; Burger 2018). A positive and increasing loop between consumption and the technosphere is therefore created. The result is an exponential escalation of technospheric throughput and associated impacts. Adhering to an outdated population narrative misdiagnoses one main cause of global environmental problems, diverts attention from consumption-driven impacts, and reinforces harmful political agendas including from xenophobia and anti-immigration rhetoric to impose reproductive control, often targeting women in low income countries of the Global South.

**Figure 1.**
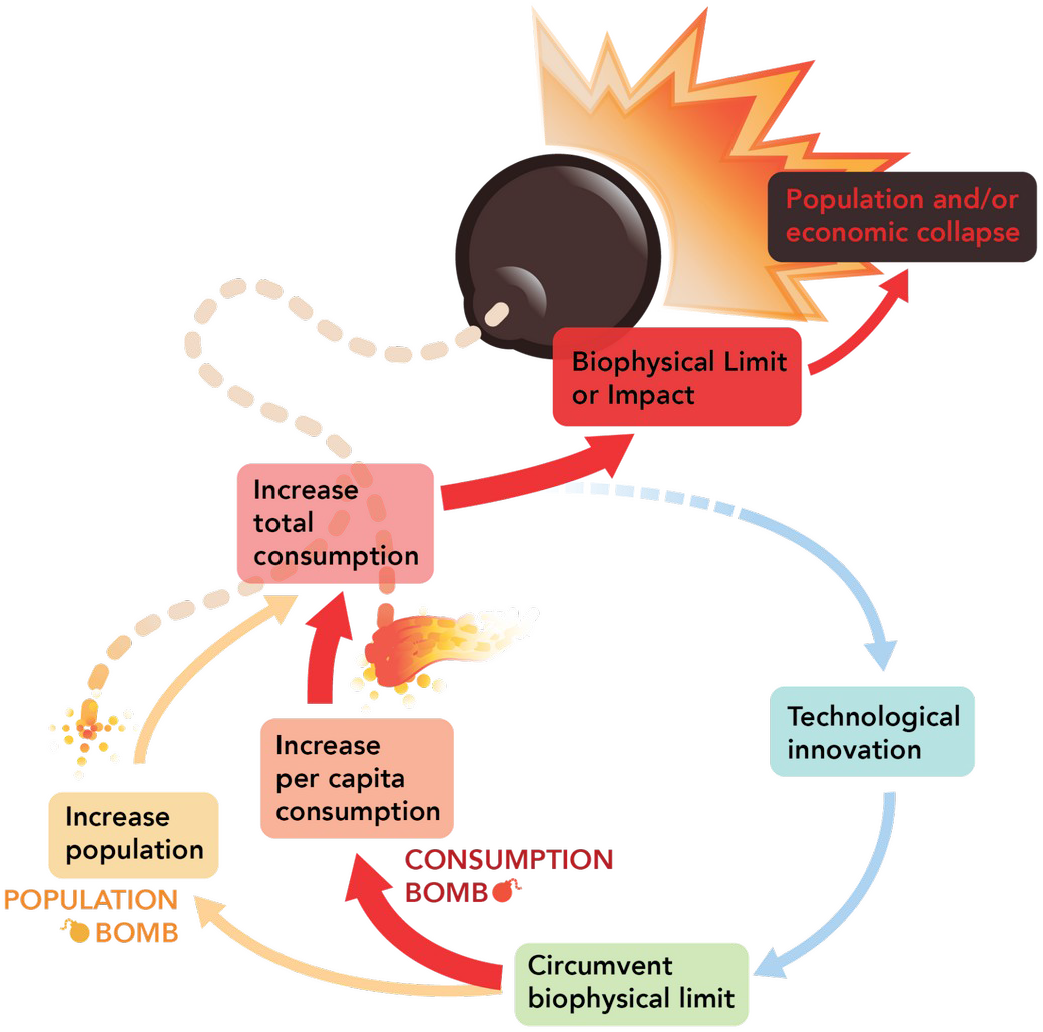
Conceptual diagram illustrating how the consumption bomb increases environmental degradation irrespective of population size. Confronted with biophysical limits or impacts, societies develop technological innovations to avert constraints and the risk of population and/or economic collapse. This enables further growth in both population and/or consumption, generating new environmental pressures that again require circumvention—thereby reinforcing the Malthusian–Darwinian dynamic. Since 1960, global population growth has slowed, yet environmental impacts have continued to rise, driven by commercial affluence and prestige-biased cultural selection. Rising per capita consumption feeds back to promote further technological evolution and expansion, producing a positive, increasing loop between consumption and the technosphere. The result is a consumption bomb: an exponential escalation of technospheric throughput and associated impacts.

In this essay, we provide cross-country analyses of global data spanning multiple decades to illustrate how population size and societal development interact to shape trajectories in energy consumption and CO_2_ production. We show that in most countries, both trends are increasing faster than population growth, signaling a growing divergence between demography and socioenvironmental impact. Our results reveal strong, persistent correlations between total and per capita energy consumption, developmental status, and CO_2_ emissions across nations and over time, indicating the limited impact of renewables in decoupling energetic demands from environmental degradation.

We introduce a simple conceptual framework to parse the contributions of population and socioeconomic development to energy demand. Our results demonstrate that the consumption bomb is fueled by countries that disproportionately contribute to global change through high and very high HDI development, driven by increments in the per capita consumption of their affluent lifestyles. This occurs despite their diminished population growth trends. Based on these findings, we flesh out the outline of a new sustainability paradigm, transcending endless growth economics and directly addressing the consumption bomb. By explicitly recognizing the biophysical limits of our planet and the necessity of consumption de-growth or zero-consumption growth (ZCG), such a paradigm would navigate a sustainable path forward for humanity.

### Energy use in human societies: the real cost of demographic transition

In less than 100,000 years, *Homo sapiens* expanded out of Africa with near-exponential growth to colonize nearly all land masses on Earth (Hamilton et al. 2009). Human societies advanced by extracting ever more resources from far corners of the Earth and transforming them into the socio-economy (Burger et al. 2012, Ellis et al. 2020, 2021, Lenton and Scheffer. 2024, Efferson et al. 2024). While hunter-gatherers use fire for heating, cooking and defense from wildlife, and employ dogs to assist in hunting and transport, they ultimately subsist on local natural resources and their metabolic demands follow the same constraints as other species (Burger et al. 2017; Burger and Fristoe 2018). Agricultural societies that emerged after the last ice age expanded energy use through plant and animal domestication, reappropriating net primary productivity into few favored species that feed or otherwise support their societies and increasing population densities (Haberl et al. 2007, 2014, Ellis 2020, 2021). With industrialization and the mass use of *extra-metabolic* energy (i.e. coal, gas and oil) since the 18th century, energy consumption has accelerated by two orders of magnitude—from ~120 watts of human biological metabolism to over 10,000 watts per person, largely from fossil fuels (Mahli 2002, Moses and Brown 2003; Hamilton et al. 2012, Burger et al. 2012, 2017, Schramski et al. 2015, Smil 2018). Modern telecoupled economies now rely on global networks of extraction, trade, communication and regulation to mobilize unprecedented flows of energy, materials, and information (Kraussmann et al. 2008, Moses 2009, Meyfroidt et al. 2010, Suweiss et al. 2011, Nesme et al. 2018).

Today, nations with high and very high development status (HDI) sustain per capita primary energy consumption ranging from 8,000 to 14,000 watts—an increase in metabolic throughput by up to 1000-fold (Fig 1B). This reliance on extra-metabolic energy in the form of fossil fuels underpins modern affluent lifestyles experienced in increasingly urban environments (Hamilton 2012, Burger et al. 2019) Although some share of this energy consumption is inherent to support social well-being indicators, such as education, life expectancy, adequate infrastructure and livelihood capacities; these nations also exhibit *conspicuous consumption*: Sisyphean pursuits of prestige, luxury, perfectionism in happiness, and commercial profit far beyond subsistence requirements (Millward-Hopkins et al. 2020).

At the same time, global fertility rates have declined from over five children per woman in 1950 to about 2.3 today, with many high-income countries even experiencing shrinking populations (Fig 2A, Lima and Berryman 2011, Barret et al. 2020, Worldometer n.d.). The demographic transition—from high to low fertility—has often been hailed as a sustainability success story. Highly developed societies require great investment to raise and educate children leading to smaller family sizes, easing future population pressures, bringing the global population closer to ZPG.

**Figure 2.**
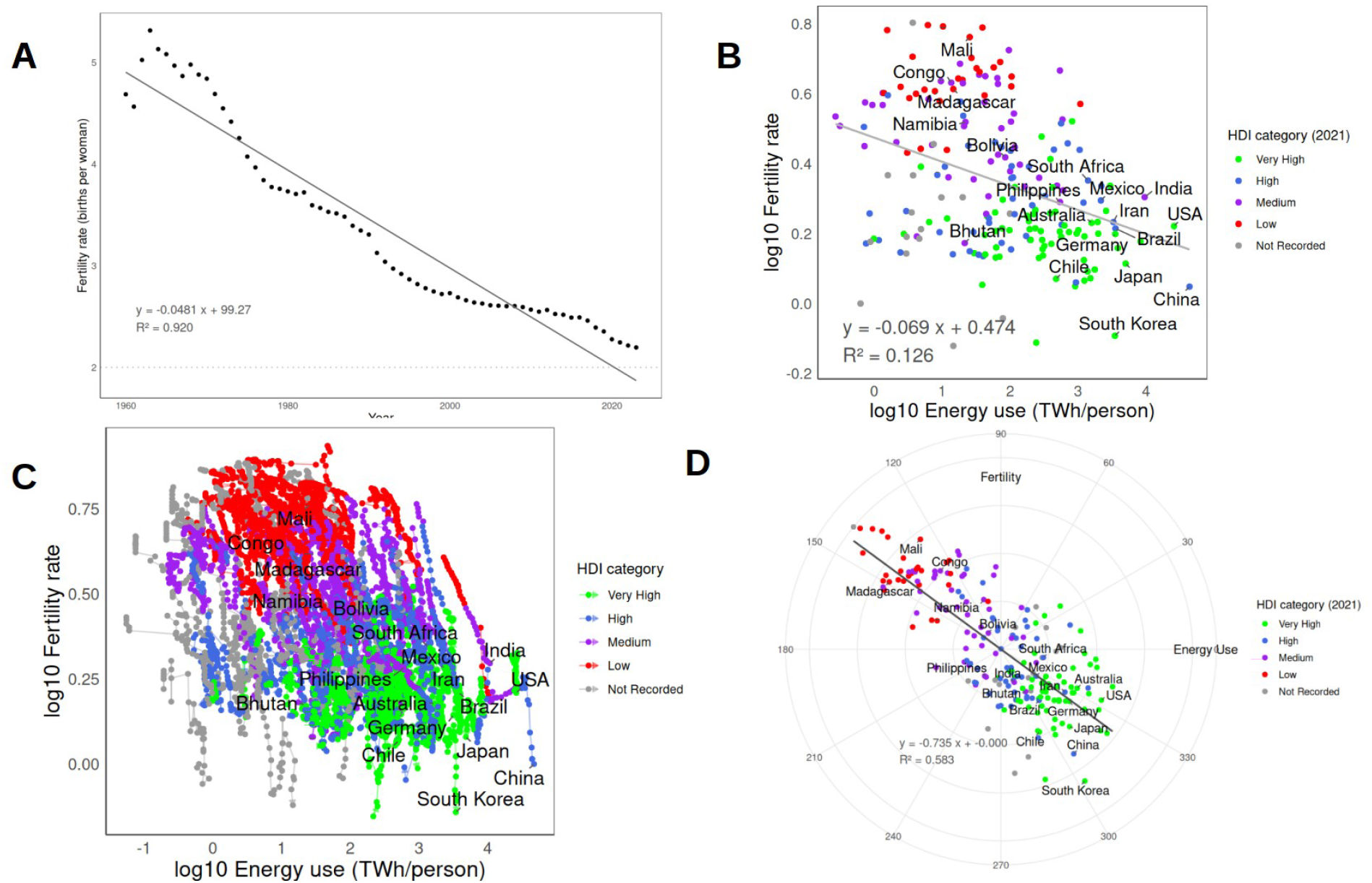
Scaling fertility and energy use across nations. **A.** Global fertility rates through time. Dashline represents non-growing population rates. **B**. Allometric relations between Total Fertility Rate vs. Primary Energy Consumption, color-coded by country HDI in 2021. **C**. Trajectories of fertility and energy consumption through time (1995–2021), color-coded by countries’ HDI. **D**. scale-adjusted national indicator (SANIs) for fertility vs. primary energy consumption allometry, scaled by population size, color-coded by countries’ HDI. The Regression line for SANI’s is shown in black.

However, the demographic transition carries steep energetic costs. Declining fertility is closely tied to the metabolic ecology and higher energy consumption in higher development nations (Fig 1B-D, Moses and Brown, 2003; DeLong et al., 2010; Brown et al. 2011, Burger et al. 2019). Countries with the lowest fertility rates, such as Japan, South Korea, and Germany, consume four to seven times more energy per person than high-fertility nations (Fig 1B,D): if fertility decline worldwide follows a similar trajectory of these high development index countries, global energy demand would be insurmountable (Delong et al. 2010).

Because over 80% of global energy still derives from fossil fuels (Fig 3), the desired demographic transition would increase contributions to greenhouse gas emissions, despite decreases to population pressure, and overwhelm efficiency gains and cleaner technologies. Indeed, countries with high and very high HDI are responsible of greater ecological footprints (Wackernagel et al. 2019) and disproportionately contribute towards exceeding planetary boundaries’ (e.g. disruption of biogeochemical flows, over-extraction of freshwater resources, contributions to emissions that drive climate change; Tian et al. 2025). The issue extends beyond emissions: even if nuclear or renewable energy provided near-limitless “clean” power, the true social and ecological costs of resource extraction, land conversion, material throughput, global commerce, and associated global change risks would remain (Farmer 2010, Lewis et al. 2017, Lyman 2021). Energy abundance would not by itself resolve the broader consumption problem.

**Figure 3.**
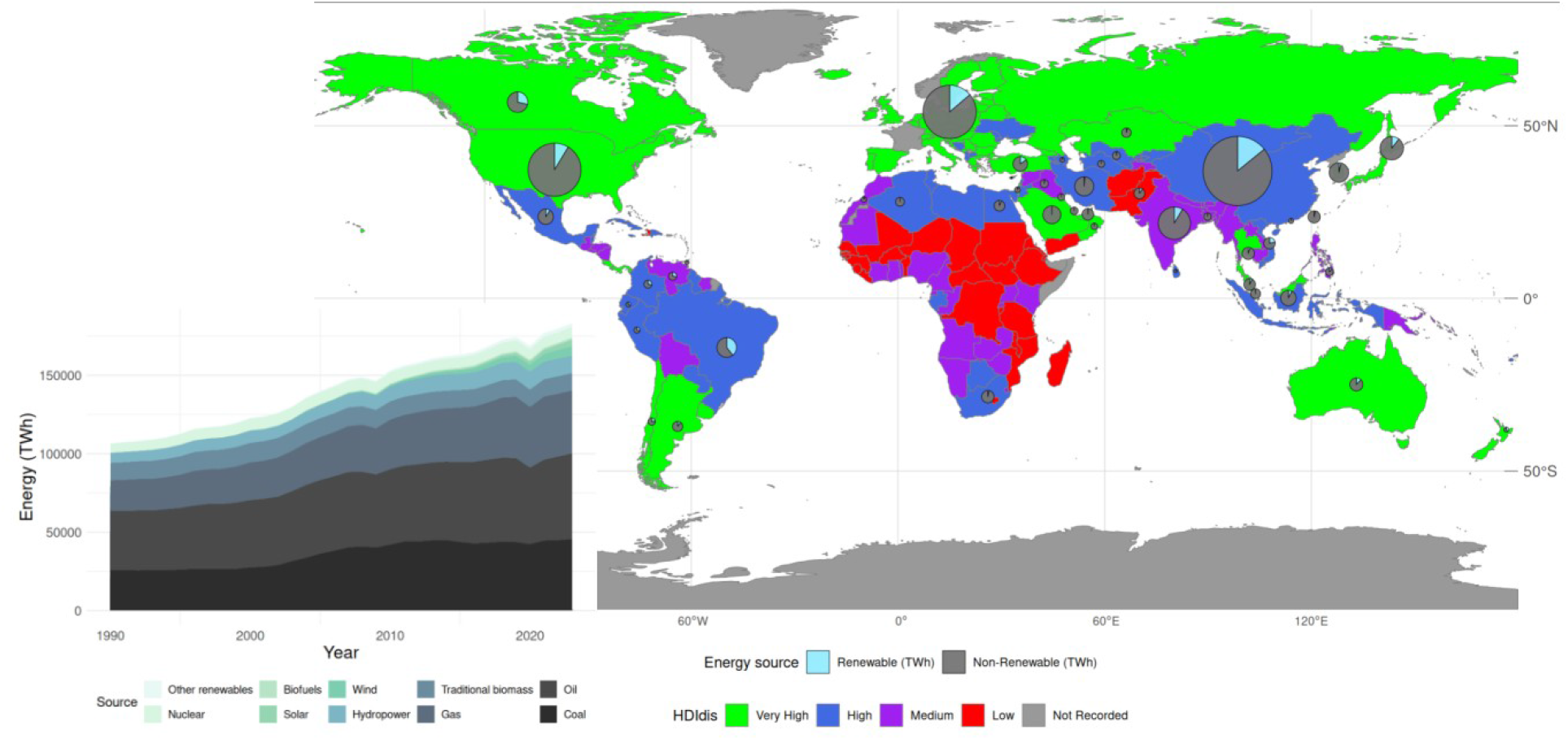
Types of energy used in modern societies. Map showing countries by Human Development Index (HDI, 2021). Donuts display the relative amount of energy consumption by country shaded by proportion from renewables versus fossil fuels. Donut size shows proportional global energy use and fraction from renewables. European countries are treated as a whole. The inset shows global energy use by type over time.

#### Box 1.

**Deep roots of the consumption bomb: overconsumption is not necessarily new**

Archaeological and historical evidence shows that the *consumption bomb*—runaway demand for resources driven by affluence, prestige, and commercial networks—long predates the industrial era. Its origins lie not in demographic pressure or basic energetic need, but in conspicuous consumption: the pursuit of prestige, luxury, and commercial profit far beyond requirements for a baseline of subsistence and wellbeing.

**Bronze Age beginnings (ca. 3000–1200 BCE)**.

The first major surge in long-distance trade emerged during the Bronze Age, when Afro-Eurasian Old World societies became linked through networks that moved metals, amber, furs, and other prestige goods across thousands of kilometers (Vandkilde 2016). This period marked the rise of urbanization, occupational specialization, and steep social inequalities—conditions that enabled elites to command surplus production and stimulate extraction in distant peripheries (Bowles and Fochesato 2024). Archaeological evidence from northern Europe suggests that even at this early stage, communities were integrated into continent-scale trade routes supplying high-value materials to Mediterranean centers (Kristiansen and Suchowska-Ducke 2015).

**Classical and Roman expansion (ca. 500 BCE–400 CE)**.

By the Roman period, resource extraction for prestige and entertainment had scaled dramatically. Amphitheater spectacles across the empire consumed vast numbers of wild animals, driving regional extirpations of elephants, lions, hippos, bears, and other large mammals in North Africa and the eastern Mediterranean (Selsvold and Webb 2020). These killings were unnecessary for survival; they served as public demonstrations of imperial wealth and power. At the same time, northern European societies—including Saami communities— participated in commercial trade systems that funneled furs, hides, and other materials toward imperial markets (Callmer et al. 2024).

**Medieval commercialization (ca. 1000–1500 CE)**.

In the late Middle Ages, intensifying trade and elite demand produced further ecological impacts. In Europe, the Eurasian beaver was hunted to near extinction for pelts and castoreum (e.g., Hoffman 2014). Similarly, wild reindeer populations were driven to local extinctions for hides and antler destined for distant markets. Reindeer mass-kill sites in the mountain region of Scandinavia show harvest levels far exceeding local dietary needs, underscoring the role of commerce and luxury demand rather than population pressure (e.g., Indrelid and Hufthammer 2011).

**Early modern globalization (ca. 1500–1900 CE)**.

The expansion of European trade companies accelerated this pattern globally. Due to increasing demand, the Russian commercial fur trade expanded in Siberia in the second half of the sixteenth century, leading to a near depletion of fur-bearing animals in roughly 150 years (Fisher 1943). The Hudson’s Bay Company and other fur-trading enterprises reshaped northern North American ecosystems (Moodie and Lehr 1981), while westward colonial expansion culminated in the industrial-scale slaughter of American bison—again, not for subsistence, but for hides, market profit, and symbolic domination of the frontier (e.g., Taylor 2011).

**A long arc toward the present**.

Across these episodes, the common driver is conspicuous consumption: resource use motivated by wealth, prestige, and expanding commercial networks. These dynamics mirror the modern *consumption bomb*, in which high-income societies operate far above basic metabolic needs, drawing on vast extra-metabolic energy flows. As cultural selection favors lifestyles of affluence and display, environmental impacts can rise even as population growth slows or reverses. The deep history of conspicuous consumption suggests that addressing today’s consumption bomb may require cultural and institutional change, not simply demographic transition or technological efficiency.

### The consumption bomb: energy use empowers affluence

While all individuals require a baseline caloric intake of about 120 watts or ~2,000 KCals per day to meet the nutritional component of decent living standards (Millward-Hopkins et al. 2020), current trend of development status (i.e. High Development Index, HDI) is a strong predictor of extra-metabolic energy use, orthogonal to population size (Fig 4A-B). Across nations, rising HDI tightly tracks increased energy consumption and therefore, CO2 emissions. This dynamic fuels the consumption bomb: a positive feedback loop where commercial affluence, urban technological lifestyles and cultural selection for prestige (Heinrich 2015, Barret et al. 2020) escalates demand for ever more resources. Even as populations stabilize at ZPG or even shrink, a condition correlated to higher developmental status, environmental impacts do not necessarily decline. Small populations with high per capita energy throughput may generate more emissions and resource depletion than larger populations living more modestly (Brown 2011).

**Figure 4.**
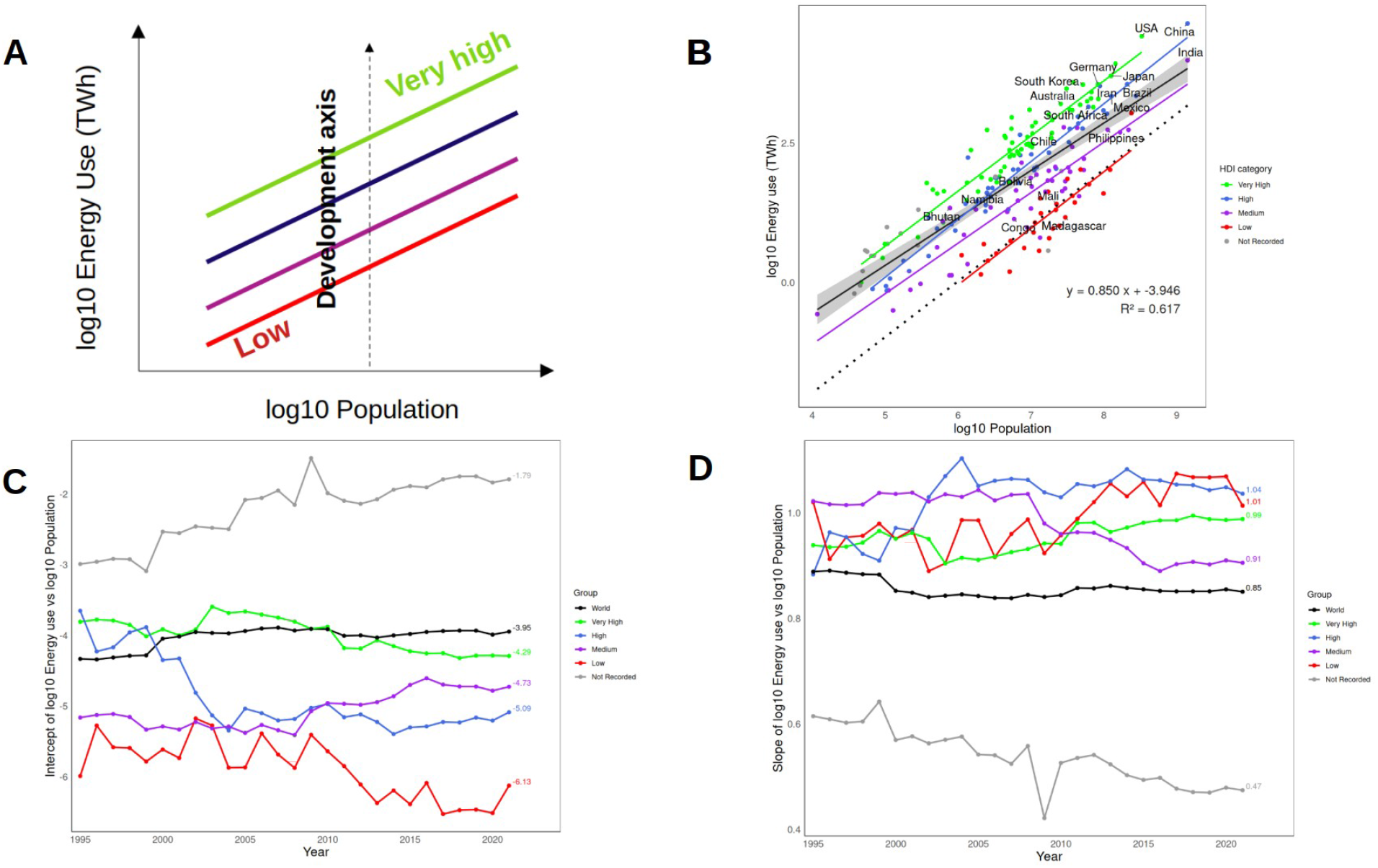
A) Conceptual diagram of orthogonal population and affluence axes B) Cross-national scaling of energy use with population size. Black dotted line represents world basal metabolic energy demand (120 TWh times total population). Dark grey line is the regression for all countries irrespective of HDI category, C) Changes in scaling prefactor (intercept) through time for each HDI category (1995–2021) D) Changes in scaling slope through time for each HDI category (1995-2021).

Increasing energy demands align with macroecological principles: as systems grow in size and complexity, they should exhibit economies of scale in response to efficiencies in distribution networks, as observed in many biological systems (West et al. 1997, Brown et al. 2004). If they, instead, require disproportionately more energy to maintain their structure and function, as has been seen for our modern telecoupled technological societies, the risk of societal collapse increases - as documented for various past human societies (Tainter 1988, Moses 2009, Brown et al, 2004, 2011; West 2017). Earlier analysis of energy consumption and population scaling showed a promising relationship of economies of scale with cities (Bettencourt et al. 2007, Bettencourt 2013). However, country level analyses that account for differences in HDI show that subliner scaling globally only arises due to the generally high population size but relatively low energy use of low development countries (Fig 4A-B, Table I).

**Table I:** Scaling indexes in Primary Energy Consumption vs Population 2021 (log scale)

|  | **slope**<br>**[95%CI]** | **Intercept [95%**<br>**CI]** | **r<sup>2</sup>** | **p-value** |
| --- | --- | --- | --- | --- |
| **All** | 0.85 [0.76,0.94] | -3.95 [-4.58,-3.31] | 0.62 | < 2.2e-16 |
| **Very High** | 0.99 [1.08, 0.91] | -4.32 [-3.75, -4.9] | 0.90 | < 2.2e-16 |
| **High** | 1.04 [1.11, 0.96] | -5.1 [-4.58, -5.62] | 0.94 | < 2.2e-16 |
| **Medium** | 0.91 [1.02, 0.79] | -4.73 [-3.92, -5.53] | 0.84 | < 2.2e-16 |
| **Low** | 1.01 [1.23, 0.8] | -6.13 [-4.56, -7.7] | 0.76 | 1.35e-9 |

Within HDI categories, energy use scales linearly with population. Moreover, their scaling relation’s prefactor (i.e. intercept) also coincides with development status (Fig 4B-C, Table I). Temporal trends show that scaling parameters tend to be consistent, at least over the past several decades (Fig 4C-D).

These results indicate that energy use is directly related to current metrics of affluence. The implications of this can be illustrated by calculating global energy increases for different hypothetical scenarios of development without increasing population: if all low HDI countries were to develop into either the medium or high HDI category, global energy would increase by only 1411.8 Twh and 12774.4 TWh, respectively, representing only 0.9% and 7.8% of global primary energy use. In contrast, if all medium HDI countries were to develop into the high HDI category, they would require 27875.6 Twh, or 17% of global energy use. Despite representing only ~35% of the global population, the greatest energy demands would occur if all the high HDI countries were to develop into the very high HDI category, requiring 46987.2 TWh of primary energy or 28.7% of the total share of world primary energy. Revisiting the developmental paradigm of modern societies becomes a necessity for sustainable outcomes.

When detailing the role of HDI development in population growth and consumption trends (Figure 5), we find that very high and high development countries are indeed the major percent of nations that decreased in population size from 1995 to 2021 (19/28 nations, 67.83%); however, only 36.84% of them (7/19 nations) were actually capable of also decreasing in energy use (Fig 5A, Table II). Most nations of the world are not only increasing in population size, but also in energy use (154/205 countries, 75.12%), with very high and high development nations representing 50% of them (77/154 countries). Actually, 63.49% of very high development nations (40/63), and 78.72% of high development nations (37/47), respectively, belong to the quadrant that increase in both population size and energy use. Clearly, energy use is empowering affluence and development through the consumption bomb.

**Figure 5.**
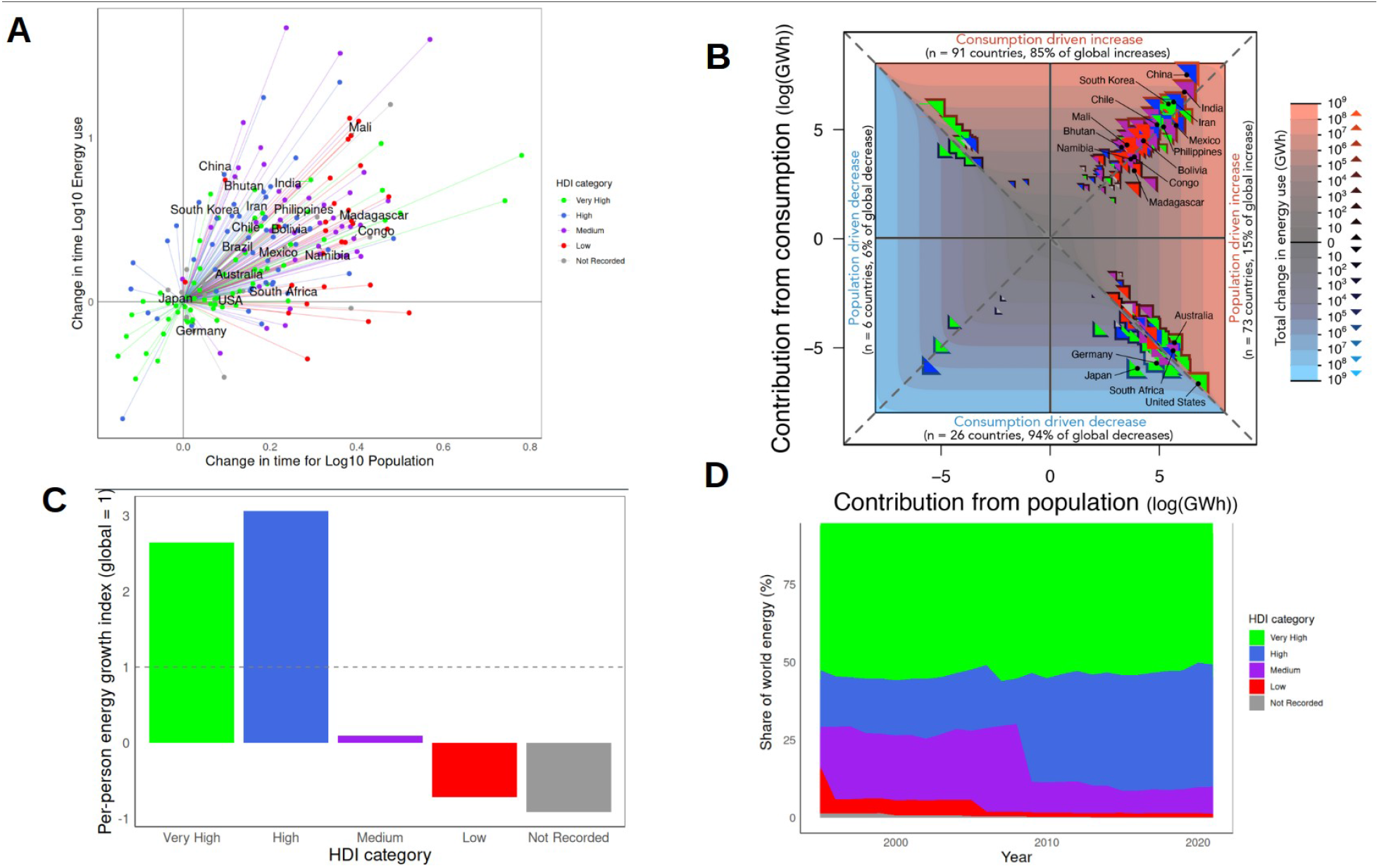
**A**. Global trends in population and energy use normalized for the same start year (1995-2021), color coded by development status. **B**. Contributions to changes in total energy use from 1995 to 2021 from population vs. per capita consumption changes. Contour lines show regions of similar total energy change from blue (decreasing) to red (increasing). Fill color for each point indicates country development status, outline color shows total energy use change over the time period, and size of points is scaled to total energy use in 1995 (log scale). Dashed lines separate the plot into four regions: increasing total energy use driven primarily by increasing per capita consumption (top in red), increasing total energy use driven primarily by increasing population (right in red), decreasing total energy use driven primarily by declining per capita consumption (bottom in blue), and decreasing total energy use driven primarily by declining population (left in blue). Inset numbers indicate the number of countries within each region (n) and their combined contribution to total change in energy use across all increasing or decreasing countries. **C**. Share of world energy use over time by HDI categories. Stacked areas show the percentage contribution of each HDI category to total global energy demand from 1995 to 2021. **D**. Per-person energy use growth index by HDI category, 1995-2021. The punctuated line at 1 represents the global-scale average per person growth in energy consumption, values above the line indicate growths larger than the world average, whereas negative values indicate declining per-capita energy demand over the period.

**Table II:** Primary Energy Consumption vs Population (1995-2021) Trajectory Counts by Quadrant (from Figure 5A).

| HDI | Energy Down,<br>Population<br>Down | Energy Up,<br>Population<br>Down | Energy Down,<br>Population Up | Energy Up,<br>Population Up | Total |
| --- | --- | --- | --- | --- | --- |
| Very High | 6 | 5 | 12 | 40 | 63 |
| High | 1 | 7 | 2 | 37 | 47 |
| Medium | 0 | 0 | 4 | 39 | 43 |
| Low | 0 | 0 | 3 | 24 | 27 |
| Not Recorded | 3 | 6 | 2 | 14 | 25 |
| **Total** | **10** | **18** | **23** | **154** | **205** |

Indeed, changes in per capita consumption, rather than changes in population, account for nearly two thirds (63%) of increasing global energy demands between 1995 and 2021. For the majority of countries with increasing energy use (55%), increases were driven primarily by increasing per capita consumption (Fig 5B), and these countries contributed disproportionately to rising global energy demand (85% of the total increases from the countries that increased energy use). Many countries that saw increasing total energy use despite declines in per capita consumption (e.g. USA and Australia) were in the very high HDI category; for these countries, energy use was so high initially, that even modest increases in population resulted in some of the greatest increases in total energy demand. Otherwise, the countries that increased their energy use the most during this time period were primarily medium to very high income countries where increases in per capita consumption outpaced increasing energy use due to growing populations (e.g. China, India, Iran, South Korea). These results illustrate the disproportionate contribution of lifestyle, rather than population pressure, in driving humanity towards planetary boundaries. However, per capita consumption was also the most important factor driving declines in total energy use. Over 70% of countries that decreased total energy use did so despite increasing populations (e.g. Germany and Japan), highlighting the crucial role of lifestyle changes in defusing the consumption bomb.

Also of interest is the role of HDI in shaping per-capita energy-demand growth over time (Fig. 5C-D). To examine this, we computed a per-person energy growth index for each HDI category (Fig. 5C), defined as the per-capita change in energy demand within each HDI category relative to the corresponding global average (global = 1; see Methods). This comparison reveals strong inequality across development levels. High HDI countries exhibit the largest increase, with per-person energy-demand growth about three times the global average, whereas the Very High-HDI group is approximately 2.5 times the global average. By contrast, the medium HDI categories remain close to zero, indicating little net per-capita growth over the study period. Even more strikingly, the low HDI category shows negative values, implying that their per-capita energy demand declined between 1995 and 2021 rather than increased.

These differences are consistent with the temporal evolution of the global energy share by HDI category (Fig. 5C). Very high HDI countries consistently account for roughly half of world energy use throughout the period, while the share of high HDI countries increases substantially over time. Together, the high and very high categories rise from roughly three-quarters of global energy use in the late 1990s to well over four-fifths by 2021. In parallel, the share associated with medium HDI countries declines markedly, suggesting that part of this shift may reflect development transitions out of the medium HDI category. However, notice that even if energy use is helping transitioning mid-developed categories, the share of energy use never diminishes for the very high ones. Low HDI countries contribute only a small fraction of total world energy use throughout the time series.

### Jevons’ paradox and the myth of decoupling

A popular belief assumes that technology will save us from economic collapse and irreparable environmental destruction–that through continual innovations, humans can develop new ideas and improvements capable of overcoming any biophysical limit they confront (Solow 1956, Boserup 1965, 1981, Borlaug 1973, Simon 1996). Indeed, through cumulative cultural evolution, humans have shown capability for ratcheting up technology into solving significant socio-environmental challenges (Boyd and Richerson 2005, Bettencourt et al. 2007, Henrich 2015), particularly those related to greater energetic and material sequestration from the biosphere toward their own societies (Ellis 2013, 2015, Snyder 2020, Kraft et al. 2021, Lenton and Scheffer 2024). However, such innovations do not intrinsically support the continuation of wealthy lifestyles, as they entail net negative externalities upon the Earth system (Meadows et al. 1972, Meadows and Randers 2012, Weinberger et al. 2017, Snyder 2020, Efferson et al. 2024). Such is the trend followed: the mass of the Technosphere exceeds all living biomass (Elhacham et al. 2020); and we have already exceeded the safe operating boundary for many different planetary limits, threatening the conditions supporting the development of human societies and the propping up of wealthy lifestyles (Brown 2011, Barnosky et al. 2012, Ellis 2013, 2015). The *cornucopian* belief of continual technological innovation and infinite growth within a finite planet is doomed at its core (Von Foerster et al. 1960, Meadows et al. 1960, Meadows and Randers 2012, Burger et al. 2012, Weinberger et al. 2017), and there is no guarantee that technological innovations and socio-economic forces will enable a self-sustaining expansion of human civilization into the hostile environment of space.

Historical trends show that whenever technology increases in efficiency (i.e. generates more energy and material per unit of energy or material required), consumption patterns do not necessarily “decouple” from biophysical processes. Instead of providing savings, introduction of more efficient technology leads to increased consumption, a phenomenon addressed as the *Jevon’s paradox*. This phenomenon has probably characterized already the earliest hunter-gatherer technologies as the energy gained from improvements in efficiency of subsistence technologies were primarily channeled to further increase foraging intensity rather than reducing the energetic costs of subsistence (Kraft et al. 2021). A striking modern example is artificial intelligence (AI), which is marketed as a tool for efficiency and optimization (Wu 2025). While AI is getting more complex and, thus, energy hungry, new models are also becoming more efficient. This makes AI cheaper and more accessible, which increases the usage of AI and the demand for massive computational resources, driving rapid growth of data centers consuming land, energy and water (e.g. Valdivia 2024, de Vries-Gao 2026). Without explicit energy caps and policy constraints, efficiency gains from AI and other digital technologies risk accelerating total consumption rather than reducing it. Efficiency alone cannot deliver sustainability; without absolute limits, technological gains are quickly undone by rebound effects.

These trends lead to a central question: can economies develop while total energy use and carbon emissions fall? Decoupling often reflects the outsourcing of carbon-intensive production to other regions, shifting rather than reducing emissions. Supply-chain accounting shows that high-income nations still drive most of global emissions through consumption, even when production occurs elsewhere (e.g. Krausmann et al. 2008, Meyfroidt et al. 2010, Suweiss et al. 2011, Krausmann et al. 2018, 2019, Wackernagel et al. 2019). Despite these complications, across countries globally, energy use and CO_2_ emissions remain tightly linked (Fig 6), exhibiting linear scaling and a prefactor also linked to affluence (Table III). These trends suggest little net gains from shifts towards renewables, even in high income countries.

**Figure 6.**
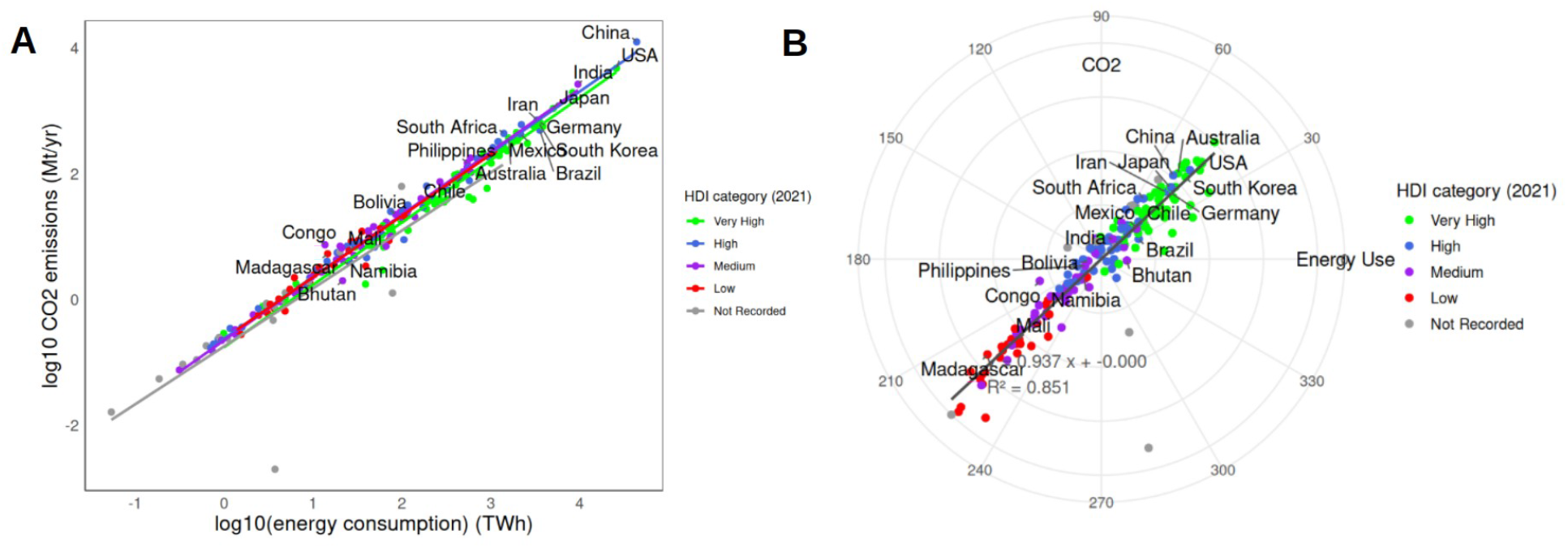
A) CO2 emissions as a function of energy consumption for countries in 2021. All slopes are linear (Table S2), suggesting limited opportunities to decouple the two. B) scale-adjusted national indicator (SANI) for CO2 vs. primary energy consumption allometry, scaled by population size, color-coded by countries’ HDI. The grey regression line indicates a positive significant relation between CO2 and energy use SANIs (i.e. nations with higher than expected energy use also produce higher than expected CO2 emissions).

**Table III:** Primary Energy Consumption vs CO2 Emissions 2021 (log scale)

|  | slope<br>[95%CI] | Intercept [95%<br>CI] | r<sup>2</sup> | p-value |
| --- | --- | --- | --- | --- |
| **Very High** | 0.96 [1.02, 0.91] | 0.84 [0.94, 0.74] | 0.96 | < 2.2e-16 |
| **High** | 1 [1.04, 0.97] | 0.65 [0.71, 0.6] | 0.99 | < 2.2e-16 |
| **Medium** | 0.98 [1.03, 0.93] | 0.68 [0.75, 0.61] | 0.98 | < 2.2e-16 |
| **Low** | 0.97 [1.04, 0.9] | 0.65 [0.71, 0.59] | 0.97 | < 2.2e-16 |

Renewable energy is essential but insufficient on its own. Current systems depend on fossil inputs, land, water, and scarce minerals. Scaling renewables to replace fossil fuels while also expanding energy access for the near billion people living in poverty (World Band, n.d.) will be impossible without major reductions in demand from high-consuming societies and redefining the multifaceted fundamentals of socioeconomic development. The illusion of “green growth” must give way to an honest reckoning with biophysical and energetic limits (Brown et al. 2011; Burger et al. 2012).

### Is a great deceleration inevitable?

All complex systems are constrained by physical limits: net primary productivity, solar energy flux, geochemical cycles, and thermodynamic boundaries. Our macroecological approach makes clear that the human system is pushing towards its own limits, primarily driven by increased resource demands and waste production by individuals in the wealthiest nations. Today’s patterns of energy use, emissions, and material extraction are overshooting these limits, destabilizing climate systems, accelerating extinction, eroding biosphere integrity, and causing considerable disruptions to Earth’s hydrosphere, atmosphere, and lithosphere.

Global energy demand has risen steadily over the past century, and despite rapid growth in renewables, the world remains overwhelmingly dependent on fossil fuels. Today, roughly three-quarters of total primary energy still comes from coal, oil, and natural gas, with the vast majority of countries consuming more nonrenewables than renewables. Fig 2 illustrates this persistent imbalance: higher HDI countries both consume more energy overall and rely heavily on fossil fuels.. These patterns further worsen the fundamental negative environmental effects of the consumption bomb: technological progress has not displaced fossil fuels; renewables only add to the total energy, supporting humanity’s trajectory of ever-rising ecological impacts.

If absolute decoupling remains elusive, then some form of *degrowth*—what systems ecologists H.T. and E.C. Odum (2006) called a “prosperous way down”—may be necessary. This does not mean universal austerity, but a deliberate, equitable contraction of energy and material use in high-consuming societies, coupled with development pathways that prioritize wellbeing and resilience over throughput and endless growth. Such a shift demands redefining prosperity in ecological, biological, and psychological terms rather than purely economic ones: sufficiency instead of accumulation, solidarity instead of competition, stability and resilience instead of perpetual growth. In this view, energy constraints are not obstacles to be engineered away, but the fundamental design parameters of a sustainable civilization.

### Toward a new sustainability paradigm

The population bomb narrative is becoming outdated. Global population trajectories are on track to reach ZPG in the coming decades. Now, the greatest ecological challenge is the consumption bomb: the accelerating use of energy and materials driven by affluence. Human macroecology provides a framework for reframing the problem around metabolism, the distribution of resources, and energetic limits. As anthropologist Leslie White (1959) others have argued (e.g. Snyder 2020, Lenton and Scheffer 2024, Ellis 2015, 2018), cultural selection for affluent socio-economic activities is a powerful and pervasive force in societies that fuels increases in per capita energy consumption. Such cultural and technological evolution has driven human expansion for millennia, fueling a Malthusian–Darwinian–Boserupian dynamic (Boserup 1965, 1981, Nekola et al. 2013; Burger et al. 2019; Cockerill et al. 2017) in which surplus energy tends to be converted into greater consumption, not restraint. Sustainability is therefore not only a technical challenge but also an evolutionary and cultural one, requiring us to break from a deeply ingrained socio-biological trajectory toward ever-higher energy flux (Richerson et al. 2024, Ellis 2024, Soogard-Jorgensen et al. 2024, Cockerill et al. 2017). Left unchecked, this dynamic will continue to drive delayed but brutal impacts on Earth’s biosphere.

Prosperity must be redefined—not as endless growth but equitable access among the population within planetary limits. Models already exist. Some real-world societies and regions show that relatively low-consumption can coexist with high-wellbeing outcomes, suggesting that sufficiency, solidarity, and social provisioning can support thriving lives. Examples include the case of Costa Rica, where life expectancies outpass those experienced in the USA, even though its income and development metrics are outperformed by the latter (Rosero-Bixby and Dow 2016); and the Indian region of Kerala, where literacy and life expectancies are among the greatest on Earth despite their low consumption (Alexander 1994).These examples highlight alternatives to industrial modernity’s high-tech cities and built environments that overly disconnect people from nature. Rebuilding relationships with the natural world—from rewilding, to preserves for public fruit foraging (Kimmerer 2021), and practising “forest bathing” (Hansen et al. 2017) —can help restore both ecological resilience and human well-being. Echoing Herman Daly’s economics of a “full world,” we must acknowledge fundamental limits to Earth and restructure our economic systems accordingly.

Defusing the consumption bomb demands transformative action. First, fossil fuel dependence must decline rapidly, alongside expanded conservation of ecosystems that safeguard biodiversity, freshwater, and carbon stocks. Second, global inequality must be confronted by redistributing energy, food, and materials more equitably, enabling low-income countries to escape ecological poverty without replicating the excesses of the affluent. Third, cities and infrastructure must be redesigned for resilience, efficiency, and local resource use. Most critically, cultural values must shift—from conspicuous consumption to sufficiency, from industrial, mono-cultive and failing distribution agriculture to a regenerative, minimal-waste generating agronomy, from fossil-fueled convenience to low-impact living, from xenophobia that incentivizes growth policies to a migration-inclusive perspective that supports elderly populations while adjusting to the new degrowth trajectory.

Developing a sustainable future within the limits of a finite planet not only depends on accounting for how many of us there are, but the resources required to support our lifestyles. Breaking the Malthusian–Darwinian–Boserupian dynamic requires a cultural, psychological, and behavioral shift away from prestige-driven consumption toward modest, resilient ways of life, supported by policy incentives. Progress must be redefined—not by the tonnes of resources extracted, but by the health and resilience of ecosystems, our communities, and the stability of our climate. The real measure of success in the Anthropocene will be our ability to turn off the engine of endless growth before it turns off the planet itself.

## METHODS

### Data

Data for total fertility rates and population numbers were obtained from the World Bank. CO2 data were obtained from the IEA-EDGAR dataset, from the 2022 European report (Cripa et al. 2022). Primary Energy use dataset comes from the U.S. Energy Information Administration (2025); Energy Institute - Statistical Review of World Energy (2025) – with major processing by Our World in Data. HDI dataset comes from UNDP, Human Development Report (2025) – with minor processing by Our World in Data. World energy use by source dataset comes from the Energy Institute - Statistical Review of World Energy (2025) and Smil (2017) – with major processing by Our World in Data. Country primary energy use dataset comes from Energy Institute - Statistical Review of World Energy (2025) – with major processing by Our World in Data.

### Analyses

All analyses were performed in R software, with codes and dataset available at github repository https://github.com/vanewi/ConsumptionBomb.

### Scale adjusted national indicators (SANIs)

Following Bettencourt et al. (2010), in order to compare urban variables between each other (energy use, CO2 and fertility rates), one must adjust for its expected scaling relation with population number *N* in the form *Y* =*Y*_0_ *N* ^*β*^, with *Y* _0_ the prefactor and *β* the scaling exponent. SANIs (*ε*) are then defined as the residuals of that scaling relation, determined as 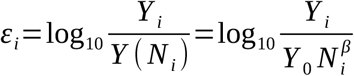, where *Y*_*i*_ is the observed value for each specific country. These SANIs are dimensionless and independent of population size. These SANIs were then normalized by the Laplace scale parameter *s*=*ε* V, estimated as the mean absolute residual of their distribution. For the polar representation, the two normalized SANIs were combined as orthogonal coordinates, with the radial distance calculated as the root of their summed squared distance 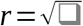 and their angle as arctan 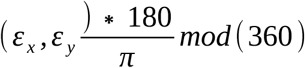.

### Per person energy growth index

To compare energy-demand growth across HDI categories, we first calculated total energy growth for each HDI category between 1995 and 2021 as *Δ E*_*class*_=*E*_(*class*,2021)_ *− E*_(*class*,1995)_. Then, we calculated the average population size of that category, as 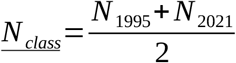, and obtained the per capita consumption of a person in that category as 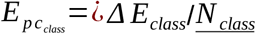. An equivalent global per capita energy demand was calculated for the world as a whole,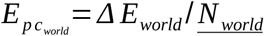, and the per person energy growth index was then defined as 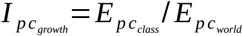, so that 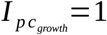 represents the global average per-person energy-demand growth, values >1 indicate above average growth, and negative values indicate declining per-capita energy demands over the study period.

## Acknowledgements

We thank many colleagues, especially Brianna Bryant, John Day, Joel Gunn, Melanie Moses, Devon Thompson, and Mathias Wakernagel, for early discussions during the development of this manuscript.

